# TransBrain: A computational framework for translating brain-wide phenotypes between humans and mice

**DOI:** 10.1101/2025.01.27.635016

**Authors:** Shangzheng Huang, Tongyu Zhang, Changsheng Dong, Yingchao Shi, Yingjie Peng, Xiya Liu, Kaixin Li, Qi Wang, Yini He, Fengqian Xiao, Xiaohan Tian, Junxing Xian, Changjiang Zhang, Qian Wu, Yijuan Zou, Long Li, Bing Liu, Xiaoqun Wang, Ang Li

## Abstract

Despite remarkable advances in whole-brain imaging technologies, the lack of quantitative approaches to bridge rodent preclinical and human studies remains a critical challenge. Here we present TransBrain, a computational framework enabling bidirectional translation of brain-wide phenotypes between humans and mice. TransBrain improves human-mouse homology mapping accuracy through: (1) a novel detached region-specific deep neural network trained on integrated multi-modal human transcriptomics to improve cortical correspondence (89.5% improvement over the original transcriptome), which revealed two evolutionarily conserved gradients explaining >50% of cortical organizational variance, and (2) random walk-based graph representation learning to construct a unified cross-species latent space incorporating anatomical hierarchies and structural connectivity. We demonstrated TransBrain’s utility through three cross-species applications: quantitative assessment of resting-state brain organizational features, inferring human cognitive functions from mouse optogenetic circuits, and translating molecular insights from mouse models to individual-level mechanisms in autism. TransBrain enables quantitative cross-species comparison and mechanistic investigation of both normal and pathological brain functions.

## Introduction

Research using human and animal models forms the cornerstone in psychiatry and cognitive neuroscience. Human studies offer direct insights into the brain function and behavior, reflecting real-world variability and clinical relevance^1^. Animal models, particularly mice, enable precise mechanistic exploration with advanced genetic tools and neural circuit manipulation, providing causal insights that are often unattainable in human research^2^. Cross-species investigation is crucial for bridging fundamental discoveries with translational applications.

Advances in whole-brain imaging technologies have created unprecedented opportunities for integrating findings across humans and mice. In humans, non-invasive neuroimaging methods, including magnetic resonance imaging (MRI), electroencephalography (EEG), and magnetoencephalography (MEG), enable researchers to study brain function and identify imaging biomarkers of psychiatric disorders^3–5^. These advances have spawned large-scale resources such as multi-site consortia (e.g., ENIGMA^6^, UK Biobank^7^) and meta-analytic platforms (e.g., Neurosynth^8^). In mice, ultra-high-field MRI, wide-field fluorescence imaging, and immunostaining of immediate-early transcription factors enable comprehensive whole-brain analysis^9,10^. However, evolutionary divergence between humans and mice, particularly in cortical regions, poses significant challenges for cross-species integration. This gap leaves the two research domains largely disconnected, limiting preclinical translation. For instance, disorders involving cortical network abnormalities, such as autism or depression, manifest as diverse biotypes in humans with distinct dysfunction patterns^11,12^. While multiple mouse models exist for these conditions^13–16^, current approaches struggle to align specific mouse models with corresponding clinical biotypes, limiting both the identification of optimal therapeutic mechanisms and efforts to enhance model validity.

Several approaches have explored human-mouse homology mapping, with greater success in evolutionarily conserved regions like subcortical nuclei^17,18^. Traditional methods rely on anatomically-defined regions of interest (ROIs), revealing homologous patterns in resting-state organization, gene effects, and sex differences^19–22^. However, this approach is limited by the subjective selection of homologous regions, leading to inconsistencies across studies. Additional challenges include inconsistent anatomical coverage (typically analyzing fewer than 10 to several dozen regions) and complex cross-species correspondence, where brain regions in one species may map to multiple regions in the other. Whole-brain spatial transcriptomics offers a more objective alternative. A pioneering study, representing the only attempt at quantitative comparison of whole-brain phenotypes, utilized transcriptional similarity based on approximately 3,000 homologous genes between mouse in situ hybridization (ISH) data and human microarray data to characterize the homology of whole-brain co-activation patterns^23^. Another study trained a supervised deep neural network on mouse ISH data to obtain latent embeddings for investigating homology and evolutionary relationships between humans and mice^24^. Recently, a self-supervised learning approach has been developed to construct a whole-brain alignment method tailored specifically to spatial transcriptomic data from both species^25^. However, these approaches show poor resolution in cortical regions, where transcriptomic and cellular heterogeneity is relatively low compared to subcortical areas^26–28^, leading these methods to primarily capture subcortical-specific features. Furthermore, purely anatomical or transcriptomic approaches overlook structural connectivity, which directly constrains functional activities and provides crucial homology insights, as demonstrated in human-macaque comparisons^29–31^.

To address these challenges, we developed TransBrain, an integrated framework for quantitative whole-brain mapping between humans and mice (**Fig.1a**). The TransBrain comprises three core steps (**Fig.1b**): spatial transcriptome matching, graph-based random walk, and mapping. In the spatial transcriptome matching phase, we employed two optimization strategies to enhance homology mapping accuracy, particularly for cortical regions. First, we integrated complementary human transcriptomic datasets, including spatially resolved microarray data^32^ and large-scale single-nucleus RNA sequencing data^28^, to improve sample size for training. Second, recognizing the distinct transcriptomic profiles between cortical and subcortical regions and the higher functional differentiation in human cortical regions, we trained a detached deep neural network model on the integrated human transcriptomic data to learn region-specific latent embeddings generalizable to mice. In the random walk phase, we constructed a heterogeneous graph comprising all brain regions from both species. Intra-species edges are defined by anatomical connectivity, derived from mouse viral tracer data^33^ and human diffusion tensor imaging (DTI) tractography. Cross-species edges were established using transcriptomic latent embeddings from the first phase, constrained by coarse-scale anatomical hierarchies. This step generates latent embeddings that integrate transcriptomic, connectivity, and anatomical hierarchical correspondence information. Finally, the mapping phase employed a dual regression approach to enable bidirectional translation of whole-brain patterns—such as imaging phenotypes—establishing a unified latent space for cross-species analysis.

**Figure 1.**
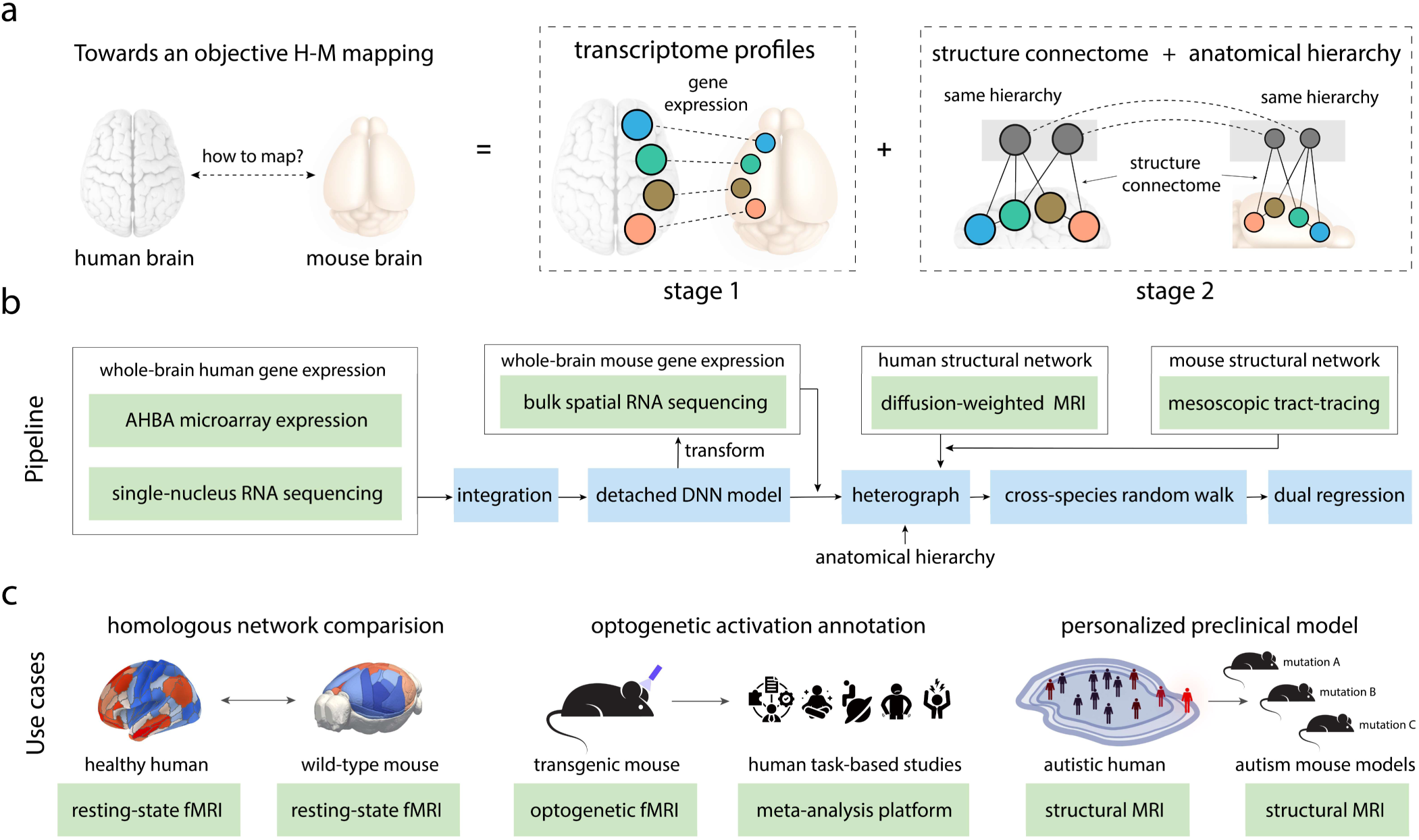
The overall framework to develop TransBrain. **(a)** TransBrain’s goal: To quantitatively map brain-wide phenotypes between humans and mice through (1) cross-species transcriptional alignment and (2) integration of anatomical hierarchies and connectivity patterns. **(b)** Overview of the TransBrain pipeline. We integrated two modalities of human brain transcriptional data: AHBA microarray expression data^32^ and single-nucleus RNA sequencing data^28^ as input. A detached deep neural network model was used to capture region-specific transcriptional embeddings for cortical and subcortical, respectively, which were then directly applied to spatial transcriptomics data of mice^40^. Using species-specific transcriptional embeddings as a bridge, we constructed a heterograph that integrates structural connectivity and anatomical hierarchies. Random walk-based graph representation learning algorithm was performed on the heterograph to obtain latent embeddings. Finally, a dual regression approach was employed to compare macroscopic imaging phenotypes between species. **(c)** Applications include: cross-species comparison of resting-state networks, translation of mouse optogenetic circuits to human brain functions, and evaluation of mouse models for human brain disorders.

To validate TransBrain’s effectiveness, we selected several classic homologous brain regions between humans and mice, particularly in cortical areas, based on a comprehensive literature review^17,18,24,25,34–36^. We assessed TransBrain’s performance by evaluating the similarity of latent embeddings in these regions. Additionally, we compared homologous relationships between cortical transcriptomic principal components, with reference to recent findings of three stable gene expression principal components in humans^37^. Our analysis revealed two highly conserved cortical transcriptomic principal components across species. Furthermore, using TransBrain, we conducted three case studies (**Fig.1c**): i) *Functional organization mapping*: We examined resting-state fMRI-based patterns in awake states, including functional networks, connectivity gradients, and whole-brain co-activation patterns, under the hypothesis that embeddings incorporating connectivity are better suited for system-level homology comparisons. ii) *Optogenetic activation mapping*: We mapped optogenetic whole-brain activation patterns of the mouse insula and dorsal raphe nucleus onto a virtual human brain and correlated these with cognitive-related maps in human standard space. This enabled plausible annotations linking these optogenetic circuits to human behaviors. iii) *Preclinical translational mapping*: We developed a normative model using multi-center autism data to capture individual gray matter volume deviation patterns, which were then mapped to various transgenic mouse models. Notably, individuals with autism showing deviation patterns significantly enriched in *magel2^-/-^*mice exhibited poorer fluid intelligence. These spatial deviation patterns also aligned closely with MAGEL2 gene expression patterns in humans. These case studies demonstrate TransBrain’s versatility in bridging fundamental neuroscience and clinical applications, from comparing brain-wide organizational features and interpreting circuit functions to advancing translational understanding of psychiatric disorders.

## Results

### Capturing latent embeddings of region-specific transcriptional profiles in humans

Beauchamp et al. has demonstrated that neural network trained on mouse brain gene expression patterns can enhance cross-species brain region matching^24^. While this approach shows good correspondence in striatal regions, it exhibits limited performance in cortical regions, particularly in supramodal subdivisions (as evident in their Fig. 5c), where the human brain exhibits substantially greater complexity in cellular diversity and functional specialization^38,39^. To improve cross-species mapping accuracy, we sought to learn region-specific transcriptional patterns directly from human brain data. A key challenge in this approach is the lack of comprehensive spatial transcriptional datasets in humans - the Allen Human Brain Atlas (AHBA) provides broad anatomical coverage but limited sampling density^32^. Importantly, a recently published whole-brain single-nucleus RNA sequencing dataset provides millions of transcriptional profiles, albeit with lower spatial resolution^28^. To leverage the complementary strengths of both resources, we developed an integration strategy by merging these datasets to achieve both high resolution and large sample size (see **Method details and Fig.2a**). PCA visualization demonstrated that merged transcriptional profiles and AHBA data points exhibit similar region-specific spatial distribution and are highly consistent in regional transcriptional patterns, indicating proper data integration. (**Supplementary Fig.1**).

**Figure 2.**
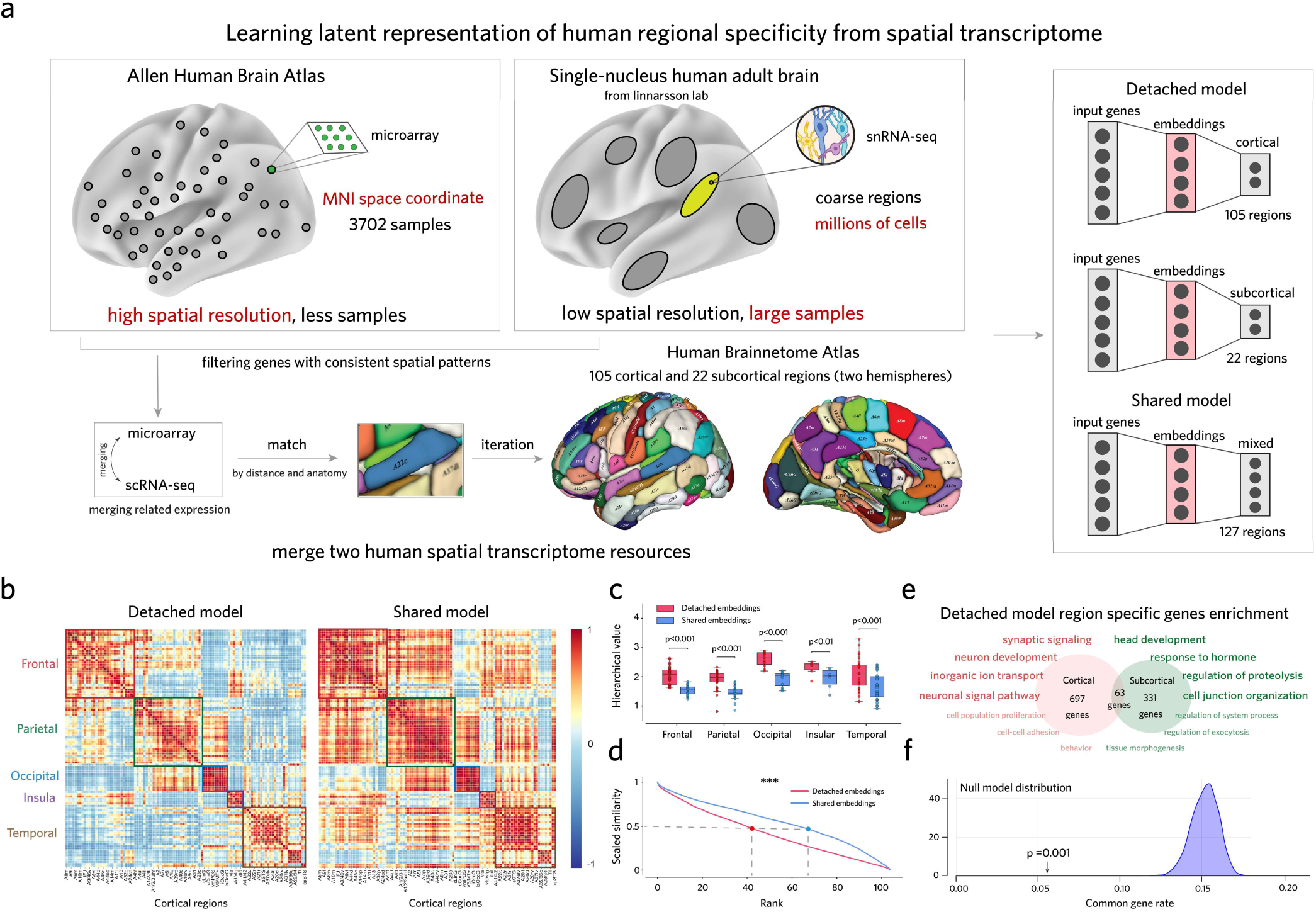
Region-specific transcriptional profiles enhanced by detached deep neural network model. **(a)** Data integration and model architecture. Left: The integration process involved selecting stable genes based on spatial expression patterns, aligning single-nucleus sampled brain regions with the Allen Human Brain Atlas, and performing a weighted combination of gene expression vectors to create a training dataset for characterizing region-specific transcriptional patterns (see **Method details**). Right: Comparison of detached model (separate cortical and subcortical processing) versus shared model (unified processing of all 127 brain regions). **(b)** Human cortical transcriptional embedding autocorrelation matrix, organized by major lobes (frontal, parietal, occipital, insular, temporal). **(c)** Hierarchical organization of transcriptional embeddings. The hierarchy value, defined as the ratio of the mean similarity of each ROI to other ROIs within the same hierarchy versus the mean similarity to ROIs outside that hierarchy in the autocorrelation matrix (detached model vs shared model; frontal lobes: *t*=10.28, *P*=3.54×10^-11^; parietal lobes: *t*=9.09, *P*=5.45×10^-10^; occipital lobes: *t*=20.77, *P*=1.49×10^-9^; insular lobes: *t*=5.32, *P*=3.15×10^-3^; temporal lobes: *t*=4.42, *P*=1.43×10^-4^; two-sided paired-samples T test). **(d)** Comparison of the specificity of transcriptional embeddings. Similarity vectors for each ROI were normalized to a range of [0,1], sorted in descending order, and averaged across all ROIs. By comparing the differences in scaled similarity at the same rank, we assessed the specificity of transcriptional embeddings (detached model vs shared model: *t*=-21.65, *P*=2.55 ×10^-40^; two-sided paired-samples T test). **(e)** GO enrichment analysis of region-specific genes derived from the detached model (cortex: 697 genes, subcortex: 331 genes, overlap: 63 genes). The cortical region-specific genes were primarily enriched in neuronal communication modules, while subcortical region-specific genes were enriched in functional regulation and homeostasis modules. **(f)** Permutation test (1000 iterations) showing minimal overlap between cortical and subcortical region-specific genes (overlap rate=0.054, *P_perm_*=0.001). \*\*\**P*<0.001.

Recognizing the substantial differences in gene expression patterns and cell-type distributions between cortical and subcortical brain regions^26–28^, we implemented two distinct training strategies using human brain region labels as learning targets. The first strategy, termed the *detached model*, trained independent modules for cortical and subcortical regions to better capture region-specific transcriptional features. The second strategy used a *shared model* that combines both cortical and subcortical data (see **Method details**). Both deep neural networks generated 500-dimensional latent embeddings in their hidden layers, which capture region-specific transcriptional patterns for subsequent analysis. Validation on strictly independent testing sets from different individuals (see **Method details** and **Supplementary Fig.2**) demonstrated that the detached models achieved significantly higher cross-subject prediction accuracy compared to the shared model (cortical; detached model: 32%; shared model: 24%; *t*=3.77, *P*=3.75×10^-4^; subcortical; detached model: 36%; shared model: 29%; *t*=2.20, *P*=0.042), with misclassifications primarily occurring between adjacent brain regions.

Analysis of the latent embedding similarity matrices revealed that the detached model captured more anatomically distinctive transcriptional patterns, particularly in cortical regions (**Fig.2b**). We quantified this improvement using two complementary metrics. First, we assessed region-specific characterization using a hierarchical value approach. We categorized the cerebral cortex into five hierarchies (frontal, parietal, occipital, insular, and temporal lobes) and subcortical regions into four hierarchies (amygdala, hippocampus, basal ganglia, and thalamus). The hierarchical value, computed as the ratio of within-hierarchy to between-hierarchy similarity, measures how well the embeddings preserve anatomical organization. The detached model showed significantly higher hierarchical values across cortical regions: frontal lobes (*t*=10.28, *P*=3.54×10^-11^), parietal lobes (*t*=9.09, *P*=5.45×10^-10^), occipital lobes (*t*=20.77, *P*=1.49×10^-9^), insular lobes (*t*=5.32, *P*=3.15×10^-3^) and temporal lobes (*t*=4.42, *P*=1.43×10^-4^) (**Fig.2c**). Second, we measured embedding specificity using a rate of decrease metric. After normalizing and sorting similarity vectors for each ROI, we found the detached model showed significantly faster similarity decay (*t*=-21.65, *P*=2.55×10^-40^; **Fig.2d**), indicating more distinct region-specific patterns. Similar improvements were observed in subcortical regions (**Supplementary Fig.3**).

The improved performance of the detached model suggested distinct region-specific transcriptional programs within cortical and subcortical areas. To validate this hypothesis, we examined gene sets that significantly contributed to model classification (see **Method details**). These genes showed significantly higher differential stability scores (DS scores) - a metric associated with diverse biological functions, brain annotations, disease relevance, and have conserved patterns in mice^27^ - compared to other genes in both cortical and subcortical regions (cortex: *t*= 32.64, *P*= 4.68×10^-210^; subcortex: *t*=3.34, *P*=8.51×10^-4^; **Supplementary Fig.4**). Gene Ontology enrichment analysis revealed distinct functional profiles: cortical region-specific genes were enriched in synaptic signaling and neuronal development pathways, while subcortical genes were enriched in hormone response and proteolytic regulation pathways (**Fig.2e**; **Supplementary Fig.5**). Permutation tests confirmed minimal overlap between cortical and subcortical region-specific gene sets (overlap rate=0.054, *P*=0.001; 1000 permutations; **Fig.2f**), indicating that the detached model effectively captured distinct transcriptional programs in these regions.

Overall, these results suggest that the detached model, which fully accounts for cortical and subcortical heterogeneity, significantly enhances the representation of transcriptional patterns in the human cerebral cortex.

### Conserved transcriptional latent embeddings between humans and mice

To explore the cross-species homology of region-specific gene expression patterns, we applied the trained models to region-averaged gene expression data from publicly available mouse spatial transcriptomics resources^40^ (**Fig.3a**, see **Method details**). Using classic human-mouse homologous brain regions based on a comprehensive literature review as benchmarks^17,18,24,25,34–36^, we quantitatively evaluated cross-species matching performance by assessing their similarity ranks among all human cortical ROIs (**Fig.3b**; **Supplementary Table.1**). The region-specific embeddings of the detached model achieved the optimal matching performance. Compared to the original transcriptome and the region-specific embeddings of shared model, the cross-species cortical homologous brain region identification ability improved by 89.5% and 84.5%, respectively. In subcortical regions, where cell-type composition and gene expression show greater heterogeneity among nuclei^26–28^, both models performed well without significant difference (**Supplementary Fig.7**). The detached model revealed consistent cross-species transcriptional similarities at finer scale, both between mouse higher-order regions (such as prelimbic) and human medial prefrontal and anterior cingulate cortex, as well as in homologous sensorimotor and visual areas (**Supplementary Fig.6c**). In contrast, the shared model failed to identify these fine-scale cortical correspondences (**Supplementary Fig.6a, b**).

**Figure 3.**
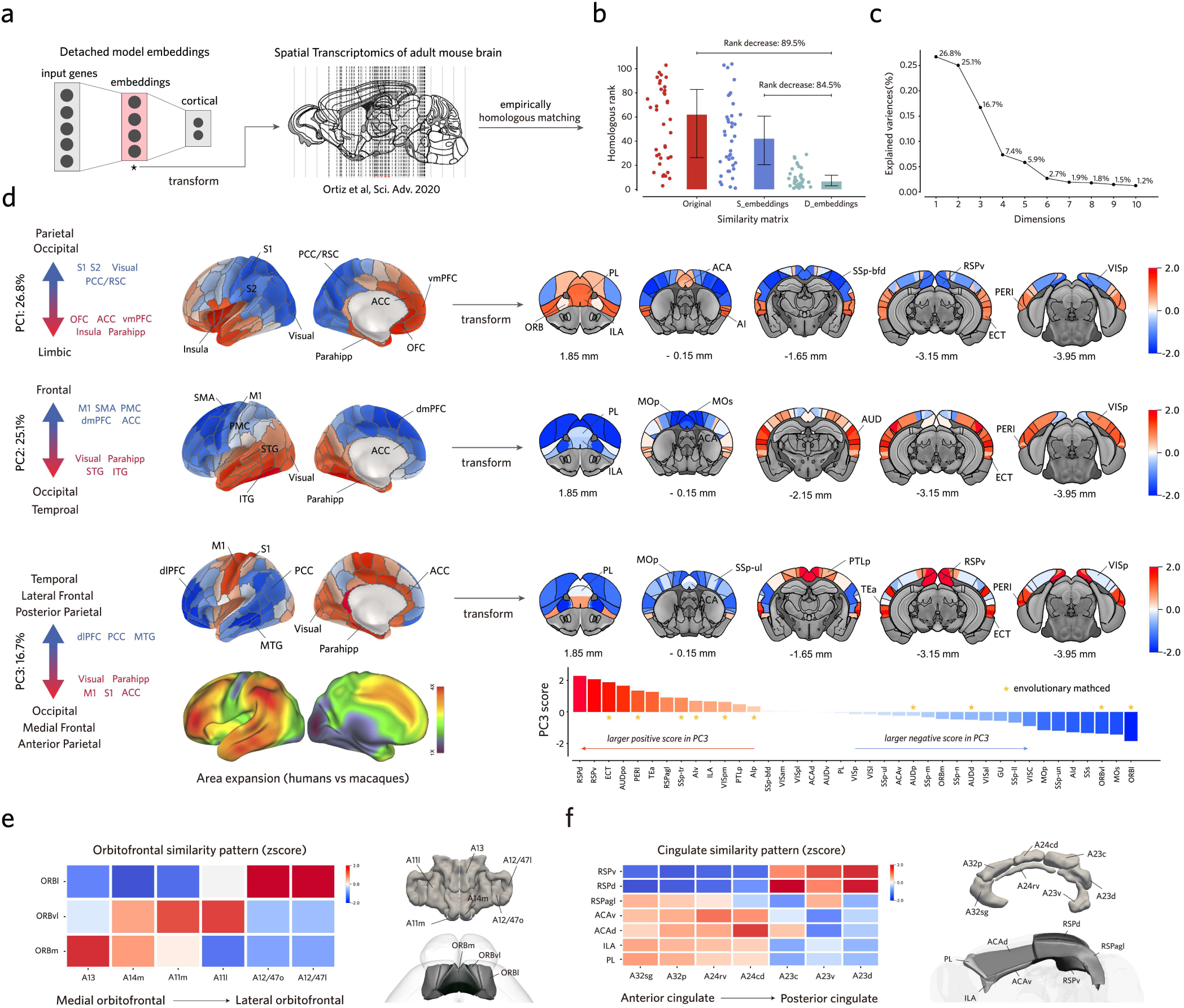
Conserved transcriptional organization patterns revealed by detached model embeddings. **(a)** Application of detached model to mouse spatial transcriptomic data^40^ for cross-species transcriptional pattern comparison. (**b**) Cross-species cortical matching performance based on cross-species similarity matrix and ranked the similarity of homologous region pairs (**Supplementary Table.1**). The detached model’s transcriptional embeddings exhibited the best cross-species matching performance. (Original: cross-species similarity matrix reconstructed from the original transcriptome; S_embeddings: cross-species similarity matrix reconstructed from the transcriptional embeddings of the shared model; D_embeddings: cross-species similarity matrix reconstructed from the transcriptional embeddings of the detached model; D_embeddings vs Original; Rank decrease: 89.5%; *t*=-8.88, *P*=1.05×10^-10^; D_embeddings vs S_embeddings; Rank decrease: 84.5%; *t*=-7.78, *P*=2.67×10^-9^; two-sided paired-samples T test). (**c**) Principal component analysis of transcriptional embeddings captured three major spatial gradients (PC1=26.8%, PC2=25.1%, PC3=16.7%). **(d)** Spatial distribution of principal components (z-scores) across species. PC1 and PC2 show conserved patterns between humans and mice, while PC3 reveals evolutionary divergence, particularly aligning with human-macaque brain expansion patterns (*asterisks indicate conserved regions). **(e)** Cross-species similarity in orbitofrontal cortex subregions follows medial-lateral gradient (z-scores). **(f)** Cross-species similarity in cingulate cortex subregions follows anterior-posterior gradient (z-scores).

Given the observed cortical correspondence, we investigated whether humans and mice share conserved transcriptional organizational patterns, as indexed by spatial gradients^41–44^. While a sensorimotor-to-association gradient has been identified in different species^45,46^, and two additional gradients were recently discovered in the human brain^47,48^, the conservation of these molecular organizational patterns between species remains unclear. Principal component analysis (PCA) of the region-specific transcriptional embeddings from our detached model revealed that the first three components explained 68.6% of the total variance (**Fig.3c**). In human cortical regions, the spatial distribution of the first three PCs closely matched the findings reported by Dear et al.^37^. Strikingly, we found high correspondence between humans and mice in PC1 and PC2 (**Fig.3d**), which together accounted for more than 50% of the total variance, suggesting substantial conservation of cortical organization between species. PC1 transitions from frontal regions (human: orbitofrontal, anterior cingulate, ventromedial prefrontal cortex; mouse: prelimbic, infralimbic, anterior cingulate) to sensorimotor and visual areas in both species. PC2 shows similar transitions from temporal/visual regions to motor/prefrontal areas across species. However, PC3 revealed species differences: while human PC3 transitions primarily from visual/sensorimotor to higher-order regions, no similar organizational pattern was observed in mice (**Fig.3d**). Notably, human PC3’s spatial distribution aligns with human-macaque brain expansion patterns^47,48^, suggesting an evolution-related transcriptional gradient. When applying this analysis to the original gene expression data, the cross-species correspondence disappeared.

We next examined two regions critical for cognitive function and psychiatric disorders: orbitofrontal and cingulate cortex. Within these regions, transcriptional similarity closely followed spatial organization: medial-lateral patterns in orbitofrontal cortex (**Fig.3e**) and anterior-posterior patterns in cingulate cortex (**Fig.3f**). Additionally, human lateral prefrontal cortex showed lower similarity with mouse higher-order regions compared to medial prefrontal cortex (**Supplementary Fig.8**), consistent with its proposed absence in mice^49^.

These results demonstrate that our detached model significantly enhanced cortical cross-species matching resolution through region-specific transcriptional embeddings, providing a molecular basis for cross-species mapping of imaging phenotypes.

### Integrating transcriptome, anatomical connectivity, and hierarchy through graph embeddings

Molecular homology between human and mouse brains has been illuminated through region-specific transcriptional profiles, but brain organization extends beyond gene expression patterns alone. The architecture of neural circuits is fundamentally shaped by anatomical connectivity patterns and hierarchical brain organization - evolutionary preserved features that govern information flow and ultimately determine brain function^31,50–53^. Cross-species comparative analyses have shown that ‘connectivity fingerprints’ effectively identify homologous regions, particularly among primates, by capturing the distinctive connection patterns that define each brain region’s functional identity^17,29,54^. To comprehensively integrate these evolutionary constraints, we developed a graph representation learning approach that leverages transcriptional similarity while incorporating both anatomical connectivity and hierarchical relationships between human and mouse brain regions (**Fig.4a**).

**Figure 4.**
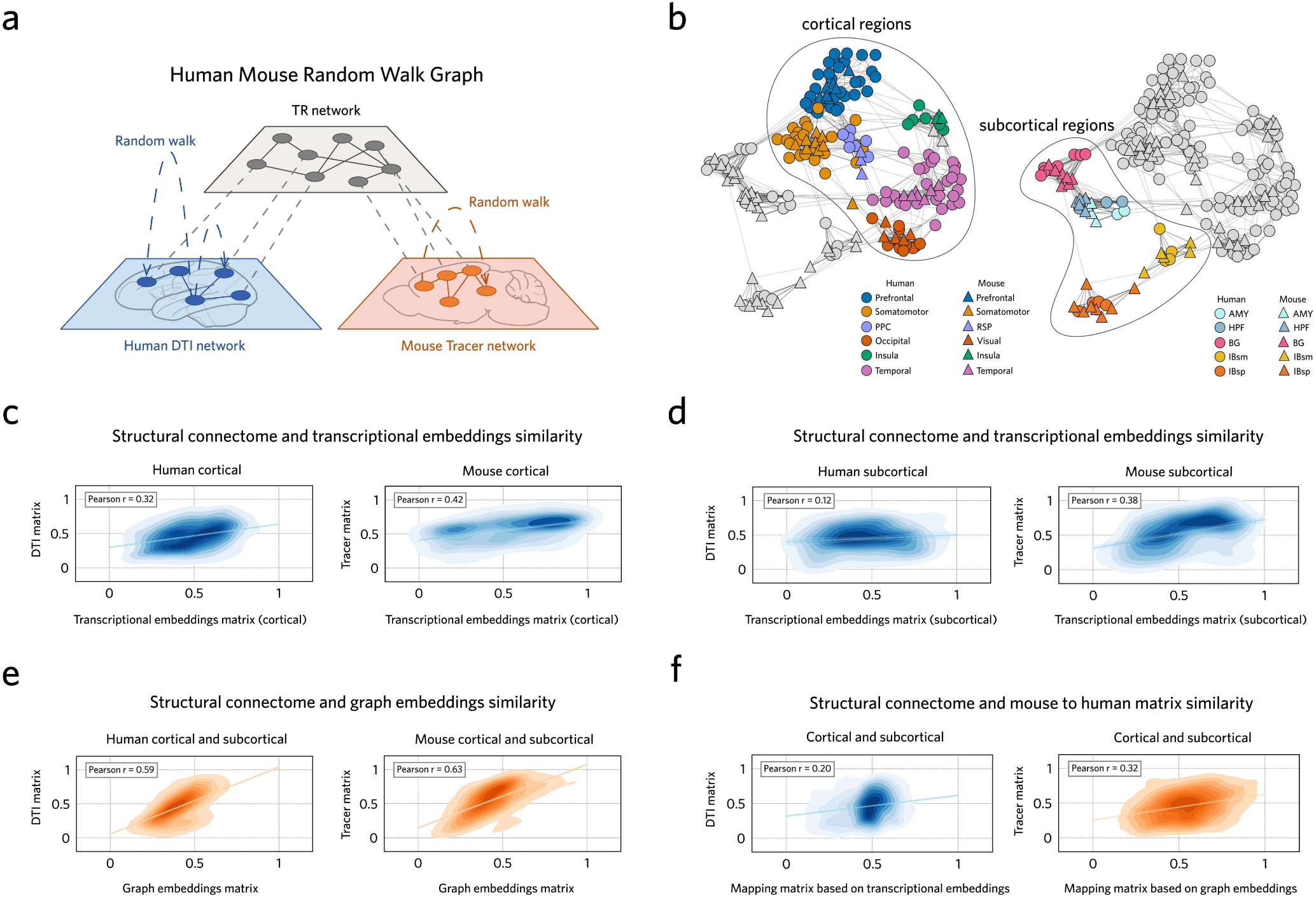
Integration of cross-species transcriptional similarity, structural connectivity, and brain hierarchies through graph embeddings. **(a)** Human-Mouse random walk process. Nodes represent brain regions connected by weighted edges based on transcriptional similarity and hierarchical constraints, with graph embeddings generated through random walk-based graph representation learning. **(b)** Two-dimensional visualization of graph embeddings: Left panel shows clustering of cortical regions, right panel highlights subcortical regions organization, both demonstrating preserved hierarchical relationships across species. **(c)** Comparison between the autocorrelation matrix reconstructed from transcriptional embeddings and the structural connectivity matrix in cortical regions (Human cortical: *r*=0.32, *P*=5.10×10^-132^; Mouse cortical: *r*=0.42, *P*=4.01×10^-29^). **(d)** Comparison between the autocorrelation matrix reconstructed from transcriptional embeddings and the structural connectivity matrix in subcortical regions (Human subcortical: *r*=0.12, *P*=0.079; Mouse subcortical: *r*=0.38, *P*=4.55×10^-15^). **(e)** Combined regions: Graph embedding autocorrelation versus structural connectivity shows improved correspondence (Human: *r*=0.59, *P=*2.23×10^-308^; Mouse: *r*=0.63, *P*=7.59×10^-240^). **(f)** Cross-species connectivity mapping: Graph embeddings outperform transcriptional embeddings in predicting human structural connectivity from mouse data (transcriptional embeddings: *r*=0.20, *P*=2.64×10^-73^; graph embeddings: *r*=0.32, *P*=2.56×10^-191^). All correlations controlled for inter-regional distance.

After the spatial transcriptome matching phase, we constructed a cross-species graph where nodes represent brain regions from both species. We weighted within-species edges using normalized anatomical connectivity derived from human DTI and mouse experimental tract-tracing data^33^. Cross-species edges were weighted by region-specific transcriptional similarities and constrained by evolutionarily related hierarchical relationships^35,36,53,55^ (**Supplementary Fig.9; Supplementary Table.2**). To ensure biological plausibility, we applied coarse-scale anatomical hierarchy constraints that only pruned cross-species connections between clearly incompatible brain divisions. Using node2vec^56,57^, we embedded this heterogeneous graph into a continuous vector space, generating latent embeddings that integrate transcriptomic, connectivity, and hierarchical information (see **Method details**). Validation analyses confirmed successful integration of both hierarchical organization and structural connectivity patterns. The embedded space revealed tight clustering of homologous regions at corresponding hierarchical levels between species (**Fig.4b**; **Supplementary Fig.11**). Quantitative assessment of the reconstructed autocorrelation matrices demonstrated substantially higher concordance with known structural connectivity patterns compared to those derived from transcriptional embeddings alone (cortical transcriptional embeddings; human: Pearson *r*=0.32, *P*=1.71×10^-132^; mouse: Pearson *r*=0.42, *P*=4.01×10^-29^; **Fig.4c**; subcortical transcriptional embeddings; human: Pearson *r*=0.12, *P*= 0.079; mouse: Pearson *r*=0.38, *P*=4.55×10^-15^; **Fig.4d**; graph embeddings; human: Pearson *r*=0.59, *P*=0.0; mouse: Pearson *r*=0.63, *P*=7.59×10^-240^; **Fig.4e**). Moreover, when we mapped mouse tract-tracing data onto human brain space, the connectivity patterns predicted by our graph embeddings showed significantly stronger correlation with actual human DTI tractography data compared to predictions from transcriptional embeddings alone (transcriptional embeddings: Pearson *r*=0.20, *P*=2.64×10^-73^; graph embeddings: Pearson *r*=0.32, *P*=2.56 ×10^-191^; **Fig.4f**).

To preserve essential transcriptional relationships while incorporating structural and hierarchical constraints, we implemented a breadth-first search strategy in the random walk process. This approach prioritized local neighborhood information, ensuring robust encoding of cross-species transcriptional similarities (see **Method details**). Comparative analysis revealed high consistency between the cross-species correlation matrices derived from graph embeddings and original transcriptional embeddings (Pearson *r*=0.53, *P*=7.79×10^-56^; **Supplementary Fig.12a**). Furthermore, both approaches demonstrated equivalent accuracy in identifying homologous regions across species (*t*=0.02, *P*=0.985; **Supplementary Fig.12b**).

### Bidirectional mapping of brain-wide phenotypes

To enable quantitative cross-species translation of brain-wide patterns, we extended our integrated embedding framework into TransBrain by implementing a dual regression approach^23^. TransBrain leverages the unified latent space - which captures region-specific transcriptional signatures, structural connectivity patterns, and hierarchical brain organization - to perform bidirectional mapping between species. This allows both the projection of mouse brain data into human neuroanatomical space and the translation of human neuroimaging features into mouse brain coordinates (see **Method details**). We demonstrate TransBrain’s versatility and translational utility through three case studies spanning different domains of neuroscience research.

### Case 1: quantitative assessment of functional network homology across species

The fundamental architecture of brain function, particularly during resting states, has been extensively characterized in both humans and mice through multiple analytical frameworks, including connectivity gradients^41,44^, temporal co-activation patterns^58,59^ (CAPs), and functional networks^58,60^. These organizational features have enhanced our understanding of fundamental brain architecture and provided valuable frameworks for investigating cognitive processes and psychiatric disorders^61^. While qualitative homology in these features has been suggested between species^62,63^, quantitative approaches for evaluating their evolutionary relationships remain limited. Here, we applied TransBrain to systematically assess the cross-species correspondence in resting-state brain organization across these functional features.

Consistent with and extending previous reports^44,62,63^, TransBrain revealed a spectrum of evolutionary conservation in functional organization between humans and mice (**Fig.5a-c**). Analysis of connectivity showed that human gradient 2 has a relatively strong correspondence with mouse gradient 2 (Pearson *r*=0.46, *P*=6.75×10^-7^; **Fig.5d** top). Human gradient 1, which captures the transition from unimodal to higher-order cognitive regions, displayed more nuanced relationships, correlating significantly with both mouse gradient 1 (Pearson *r*=0.37, *P*=8.52×10^-5^) and gradient 3 (Pearson *r*=0.53, *P*=6.56×10^-9^). While these gradients shared overall spatial patterns, regional dissociations were observed: mouse gradient 1 exhibited inverse patterns in visual areas, while gradient 3 showed reversed transitions in sensory-motor regions. Similar conservation patterns were observed in CAPs analysis, with human CAP 3 showing stronger correspondence with mouse CAP 2 (Pearson *r*=0.42, *P*=8.04×10^-6^; **Fig.5d** middle). In functional networks, we identified an unexpected correspondence between the human ventral attention network and mouse salience network (Pearson *r*=0.57, *P*=1.88×10^-10^; **Fig.5d** bottom), despite current research suggesting that mice lack established phylogenetic precursors of the VAT^42,62^. Notably, the mouse salience network also showed significant overlap with human sensory-motor regions (Pearson *r*=0.39, P=4.53×10^-5^), suggesting a more complex evolutionary trajectory. Cross-species network mappings are detailed in supplementary materials (**Supplementary Fig.13-15**).

**Figure 5.**
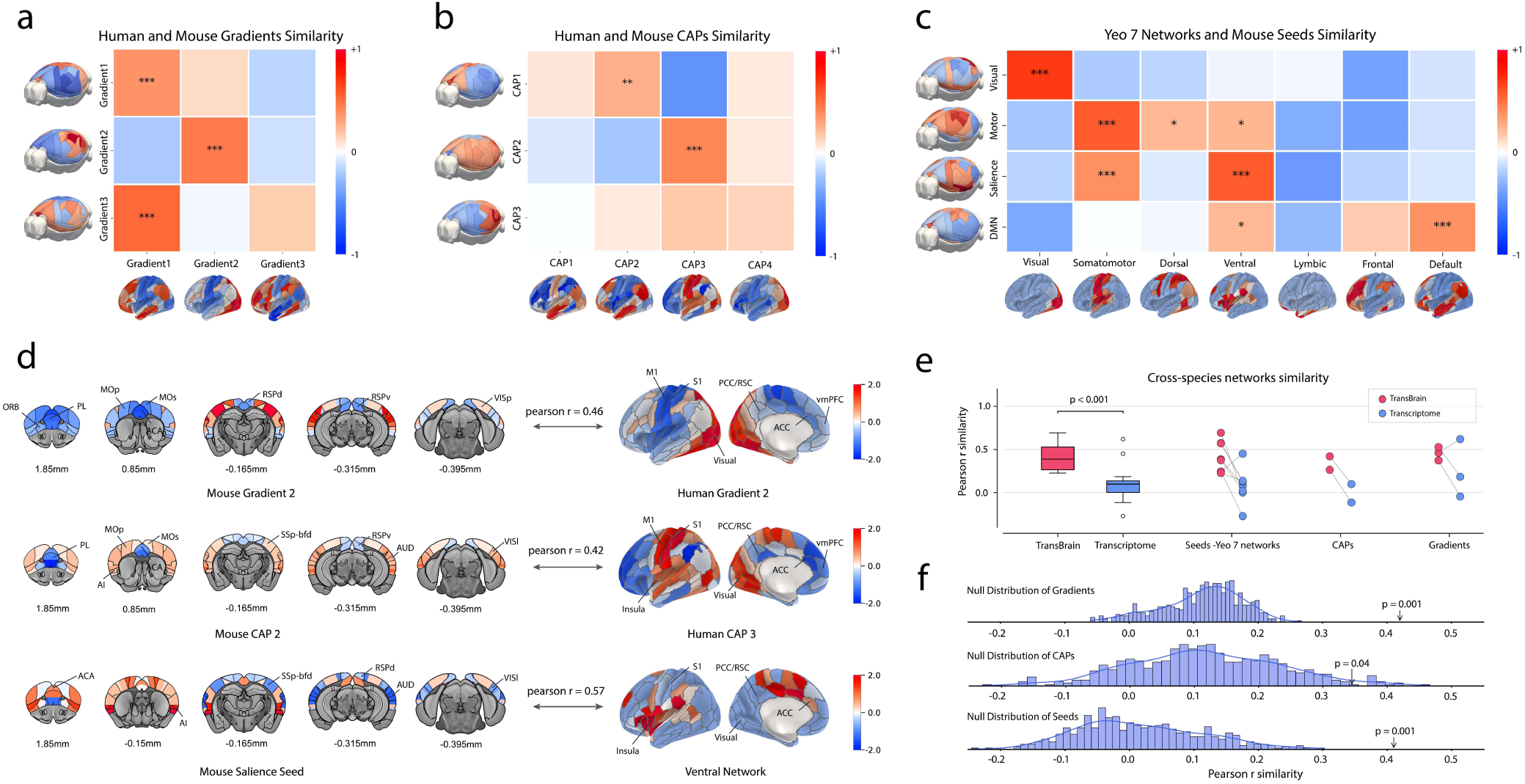
TransBrain quantitatively characterizes conserved and complex cross-species correspondences in resting-state networks. **(a**) Evolutionary relationships in gradient maps between humans and mice. (Human-Graident1 and Mouse-Gradient1; *r*=0.37, *P*=8.52×10⁻^5^; Human-Graident1 and Mouse-Gradient3; *r*=0.53, *P*=6.56×10⁻^9^; Human-Gradient2 and Mouse-Gradient2; *r*=0.46, *P*=6.75×10⁻⁷) **(b)** Evolutionary relationships in co-activation patterns (CAPs) between humans and mice (Human-CAP2 and Mouse-CAP1; *r*=0.26, *P*=6.30×10^-3^; Human-CAP3 and Mouse-CAP2; *r*=0.42, *P*=8.04×10^-6^). **(c)** Evolutionary relationships between mouse seed maps and the human yeo 7 networks (Human-Visual and Mouse-Visual; *r*=0.69, *P*=2.66×10^-16^; Human-Somatomotor and Mouse-Motor; *r*=0.57, *P*=1.55×10^-10^; Human-Somatomotor and Mouse-Salience; *r*=0.39, *P*=4.53×10^-5^; Human-Ventral and Mouse-Salience; *r*=0.57, *P*=1.88×10^-10^; Human-Default and Mouse-DMN; *r*=0.38, *P*=7.72×10^-5^). **(d)** Visualization of the evolutionary matching anatomical details between the human and mouse unimodal networks, as well as the evolutionary matching anatomical details between the mouse salience seed map and human ventral network. **(e)** Using the evolutionarily significant matching (p < 0.05) cross-species network correlation values (Pearson *r*) from TransBrain as the baseline, we compared the evolutionary matching results based solely on transcriptome, TransBrain exhibited significantly higher matching similarity between species (TransBrain vs transcriptome; *t*=4.58, *P*=6.30×10^-4^; two-sided paired-samples T test). **(f)** The null model, created by randomly shuffling the structural connectivity of humans and mice while maintaining node degree and other parameters, was used to evaluate the necessity of incorporating structural connectivity information. The mean of Pearson *r* correlation coefficient of significantly matched network organization maps, based on TransBrain, served as the baseline for comparison (Gradients; baseline=0.42, *P_null_*=0.001; CAPs; baseline=0.35, *P_null_*=0.04; Seeds-yeo 7 networks; baseline=0.41, *P_null_*=0.001; 1000 repetitions). \**P*<0.05, \*\**P*<0.01, \*\*\**P*<0.001.

To quantitatively assess the performance gains of our integrated framework, we next compared TransBrain against mapping based solely on region-specific transcriptional embeddings. Using the Pearson correlation coefficients of the significantly matched network organization maps as a baseline, TransBrain achieved substantially higher cross-species matching similarity (*t*=4.58, *P*=6.30×10^-4^; **Fig.5e**), with consistent improvements across diverse functional organizational features. To control for potential selection bias in TransBrain-identified network matches, we conducted parallel analyses using transcriptional-only mapping, which revealed several biologically implausible correspondences (**Supplementary Fig.16**), including spurious matches between mouse sensory-motor and human visual networks. To rigorously evaluate the contribution of structural connectivity constraints, we implemented a density and degree-preserved null model with randomized connectivity^64^. Although this null model maintained transcriptional relationships (Pearson *r*=0.68, *P*=9.33×10^-115^; **Supplementary Fig.17a**), it exhibited significantly diminished performance in cross-species network organization mapping (connectivity gradients: *P*_null_=0.001; CAPs: *P*_null_=0.04; functional networks: *P*_null_=0.001; 1000 repetitions, **Fig.5f**). These systematic validations establish that accurate cross-species mapping of functional brain organization requires the integration of multiple evolutionary constraints - transcriptional profiles, structural connectivity, and hierarchical relationships.

### Case2: Translation of mouse optogenetic circuits to human brain functions

Optogenetic fMRI provides detailed maps of whole-brain responses to circuit-specific manipulations in mice^65^, yet translating these findings into human behavioral contexts remains challenging. Conversely, human task-fMRI has identified robust task-related activation patterns linked to specific cognitive and behavioral states^66^, but lacks causal circuit-level evidence. Here, we leveraged TransBrain to bridge this translational gap by mapping optogenetically-driven circuit patterns into human brain space and correlating them with established task-fMRI databases^8^.

TransBrain predicted distinct human brain activation networks for both insula and dorsal raphe nucleus (DRN) stimulation. The predicted insula stimulation network encompassed insula, thalamus, ventromedial prefrontal cortex and lateral orbitofrontal cortex (**Fig.6a**). This network architecture, centered on thalamus relay circuits, suggests coordinated integration of sensory processing and emotional regulation^67^. The predicted DRN stimulation network revealed a broader activation across emotion and cognitive control regions, including anterior cingulate, ventromedial and dorsolateral prefrontal cortices, posterior cingulate, insula, medial orbitofrontal cortex, amygdala, and striatum^68–70^(**Fig.6b**).

**Figure 6.**
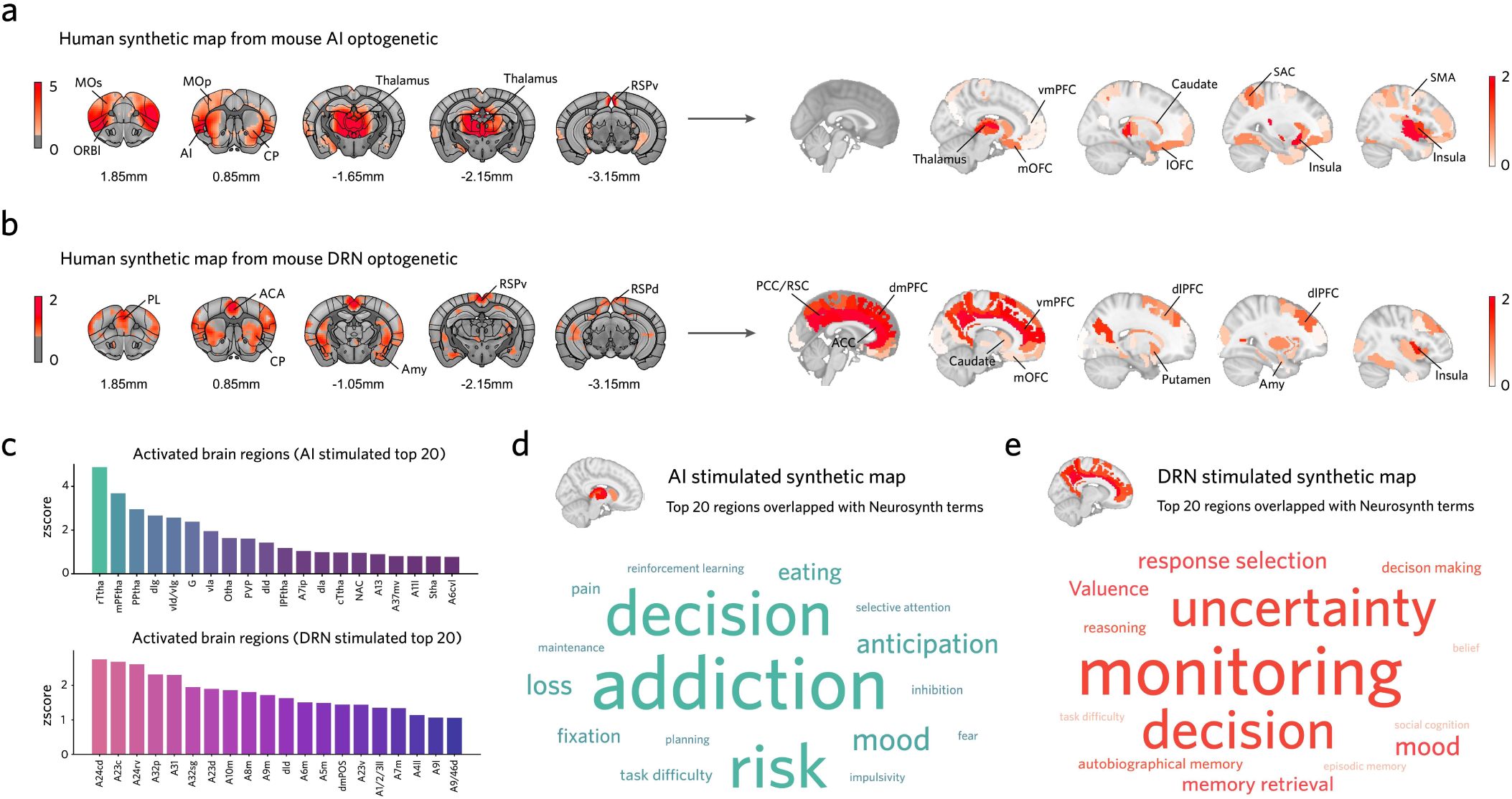
TransBrain maps mouse optogenetic circuits to human behavior-related brain patterns. **(a)** Mouse insula optogenetic activation patterns mapped to predicted human brain networks (positive activations only, z-score>0). **(b)** Mouse dorsal raphe nucleus optogenetic activation patterns mapped to predicted human brain networks (positive activations only, z-score>0). **(c)** Anatomical distribution of top 20 regions in both predicted networks. **(d)** Neurosynth behavioral mapping of predicted human insula network (top 20 regions) reveals associations with decision-making, addiction, anticipation, risk assessment, and mood regulation. **(e)** Neurosynth behavioral mapping of predicted human dorsal raphe network (top 20 regions) reveals associations with uncertainty processing, cognitive monitoring, decision-making, and valence processing.

To explore the behavioral relevance of these optogenetically-driven networks, we quantitatively compared the top 20 predicted activation regions from each stimulation condition (**Fig.6c**) against human task-fMRI meta-analysis maps (see **Method details**). Insula-driven circuit predictions aligned with decision-making, addiction, anticipation, risk assessment, mood regulation, and pain processing tasks (**Fig.6d**). DRN-driven predictions corresponded with cognitive monitoring, decision-making, valence processing, and mood regulation (**Fig.6e**). These examples demonstrate that TransBrain can translate mouse circuit manipulations into human-relevant predictions, offering a novel computational avenue to link optogenetic findings with established human cognitive maps.

### Case3: Linked gene mutations to imaging phenotype deviations in autism

Understanding the molecular mechanisms underlying individual heterogeneity in psychiatric disorders remains a fundamental challenge in translational neuroscience. While mouse models enable direct investigation of gene-phenotype relationships, translating these findings to human clinical presentations has been limited by the lack of quantitative cross-species comparison frameworks^2^. Here, we demonstrate TransBrain’s utility in bridging this gap using autism as a proof-of-concept, establishing an objective approach to link human imaging phenotypes with genetic mouse models.

To address the considerable heterogeneity in autism, we implemented a normative modeling approach^71^ - which characterizes atypical brain features by comparing each individual against a statistical model of typical brain development - to quantify region-specific brain volume deviations in patients. In parallel, we quantified mutation-specific structural alterations in mouse models by computing standardized volume differences (Cohen’s d) between mutant and control groups (see **Method details**). Using TransBrain, we mapped these mouse structural alterations into human brain space to compare with individual patient abnormalities. For each patient, we calculated risk scores that quantify how well their brain volume deviation patterns match those predicted from different mouse genetic models (**Fig.7a**). Additionally, we computed “Allen expression-based scores” by correlating patient brain abnormalities with the spatial expression patterns of corresponding mutated genes from the Allen Human Brain^32,72,73^. The significant correlation between these two independent scoring approaches (Pearson *r*=0.20, *P*=9.65×10^-12^; **Fig.7b**), suggests that TransBrain can potentially link clinical heterogeneity with specific genetic mechanisms.

**Figure 7.**
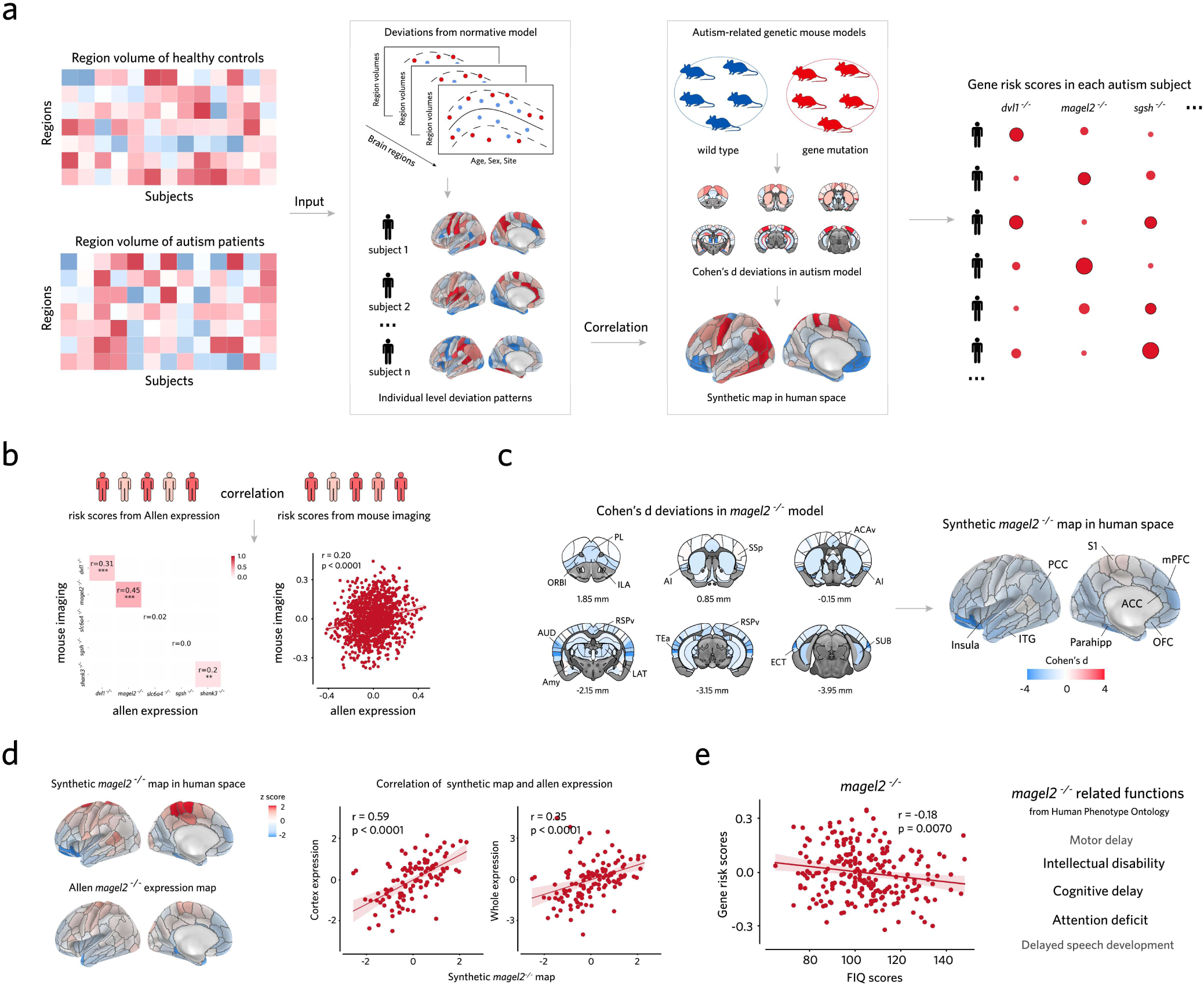
TransBrain enables assessment of autism risk mechanisms across species. **(a)** Generation of mouse imaging-based risk scores: TransBrain maps group-level structural differences from mutant mouse models to human brain space, then compares with individual autism patients’ deviations (calculated using a normative model based on healthy controls) to generate personalized risk scores. **(b)** Validation of risk scores: significant correlation between mouse imaging-derived and Allen gene expression-based risk scores (*r*=0.20, *P*=9.65×10⁻¹²). **(c)** Brain regions with significant volume deviations in the *magel2^-/-^* mouse model and synthetic deviations map in human space from TransBrain. **(d)** *Magel2^-/-^* model’s predicted human patterns significantly match MAGEL2 gene expression distribution (whole brain: *r*=0.35, *P*=4.78×10^-5^; cortex: *r*=0.59, *P*=2.73×10^-11^). **(e)** Clinical relevance: risk scores based on the *magel2^-/-^*mouse model was significantly negatively correlated with FIQ in autism patients (*r*=-0.18, *P*=0.007).

Among all examined mouse models, *magel2-/-* showed the strongest agreement between imaging-based and expression-based scores (Pearson *r*=0.45, *P*=3.01×10^-13^), followed by *dvl1-/-* (Pearson *r*=0.31, *P*=1.29×10^-6^) and *shank3-/-* (Pearson *r*=0.20, *P*=2.14×10^-3^) (**Fig.7b**). Furthermore, when analyzing spatial correspondence between mouse imaging alterations and human gene expression patterns, the *magel2-/-* model showed significant alignment with MAGEL2 gene distribution (Pearson *r*=0.35, *P*=4.78×10^-5^; **Fig.7d**).

Given these compelling cross-species alignments, we further investigated the *magel2-/-* model’s translational relevance through anatomical and behavioral analyses. In mice, significant volumetric alterations were observed in prelimbic, infralimbic, anterior cingulate, insula, somatosensory, ventral retrosplenial, and ectorhinal regions. When mapped to human space, these corresponded to anterior cingulate cortex, medial prefrontal cortex, orbitofrontal cortex, insula, and primary somatosensory cortex (**Fig.7c**) - regions involved in decision-making, cognitive control, and somatosensory processing^67,68,74,75^. Previous studies have identified magel2-/-associated phenotypes including cognitive delays and intellectual disability^76–78^. Consistently, we found that magel2-/- based risk scores negatively correlated with fluid intelligence quotient (FIQ) in autism patients (Pearson r=-0.18, P=0.0070; **Fig.7e**), suggesting conserved gene-brain-behavior relationships across species.

These findings suggest TransBrain’s potential for quantitatively assessing cross-species correspondence in brain alterations. As standardized multi-modal MRI datasets of mouse psychiatric disease models become available, this framework could facilitate mechanism-oriented investigation in psychiatric disorders through systematic cross-species comparison.

## Discussion

Recent advances in whole-brain imaging technologies have generated unprecedented amounts of data across humans and mice, offering new opportunities for cross-species integration. However, translating these findings between species remains challenging due to fundamental differences in brain organization. In this study, we developed TransBrain, a framework to quantitatively translate whole-brain phenotypes between humans and mice. At the methodological level, the framework addresses two fundamental challenges in cross-species translation: i) we enhanced region-specific transcriptional matching between species by integrating complementary human transcriptomic datasets (spatially-resolved microarray data^32^ and large-scale single-nucleus RNA sequencing data^28^) and implementing a detached supervised learning approach, achieving substantially improved cortical correspondence; ii) we established a unified brain latent space by integrating cross-species data modalities from both humans and mice, including transcriptional similarity, anatomical hierarchies, and structural connectivity (human DTI tractography and mouse viral tracing^33^), through random walk-based graph representation learning. Combined with a dual regression approach, this unified space enables bidirectional transformation of brain-wide phenotypes between species. Through comprehensive validation and three case studies - functional network homology assessment, optogenetic circuit translation, and autism model evaluation - we demonstrated how TransBrain enables systematic analysis of evolutionary conservation and preclinical translation.

The field of cross-species brain mapping faces unprecedented obstacles when translating between humans and mice. These difficulties, which far exceed those encountered in human-primate comparisons, arise from extensive cortical expansion and reorganization in primates^79–81^. Previous research has identified multiple conserved features between mouse and human brains, including cell composition^38,82–84^, gene expression patterns^24,25,27^, and connectivity fingerprints^17,54,85^. These conserved features suggest fundamental constraints in brain organization across species, reflecting intrinsic relationships between molecular patterns and region-specific functions.

While improving the resolution of transcriptional matching between species, particularly in cortical regions, remains a critical limitation, we implemented several methodological improvements to address this. First, we integrated complementary human transcriptomic datasets to leverage both the broad anatomical coverage of microarray data^32^ and the large-scale sampling of single-nucleus RNA sequencing^28^. Second, recognizing that human cortex exhibits more refined molecular and functional specialization than mice^86^, we trained our model on human data to learn region-specific features. Furthermore, our separate training strategies for cortical and subcortical regions enhanced the detection of regional transcriptional signatures. Surprisingly, our analysis revealed that the first two principal components of cortical transcriptional patterns, explaining over 50% of the variance, are highly conserved between species, suggesting substantial conservation of spatial molecular organization. The third component, however, aligns with evolutionary expansion, corresponding to higher-order networks in humans. Notably, we also observed conserved intra-regional transcriptional organization between species, particularly in key cognitive areas such as the orbitofrontal cortex (medial-lateral gradient) and cingulate cortex (anterior-posterior gradient), further supporting the evolutionary preservation of fundamental molecular-spatial patterns.

Although transcriptional similarity-based matching provides a foundational correspondence for cross-species brain region relationships, it has inherent limitations in cross-species mapping, such as producing one-to-many relationships. While some of these relationships can be biologically meaningful within the same hierarchical level (e.g., mouse prelimbic cortex corresponding to both human medial prefrontal cortex and anterior cingulate cortex), others can produce spurious matches across different hierarchical levels. For instance, regions may share certain degrees of transcriptional features despite clear evolutionary divergence (e.g., human medial prefrontal cortex showing moderate similarity with mouse visual areas), potentially introducing errors in phenotype mapping. Building on insights from a human-macaque comparative study^30^ that successfully integrated regional and connectivity information, we recognized the importance of extending beyond transcriptional patterns alone. Brain function is fundamentally constrained by structural connectivity, which shapes both functional activity and the propagation of disease-related alterations^87–89^. These structural constraints provide essential topological boundaries for cross-species translation by guiding regional interactions and influencing global brain organization. To leverage these organizing principles, we developed a cross-species random walk-based graph representation learning method that integrates hierarchical constraints and species-specific structural connectivity while preserving transcriptional relationships. Our validation analyses demonstrated that this integrated approach significantly enhanced the cross-species correspondence of resting-state functional organization patterns, supporting its biological relevance.

While our framework enables quantitative comparisons between humans and mice, several limitations warrant consideration. First, TransBrain was designed for whole-brain phenotype translation, potentially overlooking specific regional homologies. Therefore, when evaluating mouse models with strong regional priors, complementary region-focused methods could be valuable for comprehensive assessment. Second, given the substantial evolutionary gap between humans and mice, significant correlations identified by TransBrain should be interpreted as evidence for similar organizational patterns rather than definitive evolutionary homology. Third, current applications in both human and mouse datasets have not reached their full potential. As spatial transcriptomics and connectomics technologies advance and more data becomes available^26,90,91,91–94^, the mapping accuracy of TransBrain will continue to improve. Future research should also integrate evidence from emerging sequencing technologies for more comprehensive cross-species mapping.

Moreover, advancements in high-resolution whole-brain imaging techniques in mice, the establishment of open data resource platforms, and the development of standardized public protocols will greatly facilitate cross-species translational research between humans and mice. For instance, in **Case 1**, the mouse brain networks we characterized were derived from awake-state fMRI data acquired at a single center. It is therefore essential to evaluate the reproducibility of these brain networks across different research centers and to consider the influence of data acquisition strategies and post-processing protocols. Excitingly, efforts in this direction are already underway^95–97^. In **Case 2**, we used TransBrain to obtain whole-brain activation patterns of the insula and dorsal raphe nucleus mapped into human space. However, achieving finer-scale mapping of brain regions and cell-type-specific regulation will provide a more detailed perspective for understanding the principles underlying brain circuit functions^65^. Similarly, in **Case 3**, we presented a conceptual example to evaluate the correspondence between specific genetic mouse models and human imaging phenotypes, based solely on brain structural biases. The integration of large-scale, multimodal mouse model data holds great promise for transforming this concept into a more objective reality, thereby deepening our understanding of the complex molecular mechanisms underlying psychiatric disorders. In future, as large-scale mouse imaging databases emerge, TransBrain could transform cross-species comparative studies by enabling joint analysis of mouse and human phenotypes on a unified scale. This framework promises to advance both mechanistic understanding in translational neuroscience and therapeutic development through improved animal model evaluation and biological mechanism discovery.

## Methods

### Human atlas

All cortical regions were derived from the Human Brainnetome Atlas, which is constructed based on whole-brain structural connectivity^98^. For subcortical structures, we adopted a hybrid approach: the amygdala and thalamus were parcellated using the Human Brainnetome Atlas, while the hippocampal and basal ganglia were parcellated using a manually delineated atlas based on T1 and T2 weighted images from 5 healthy subjects^99^ and 3D Allen Brain Atlas^100^, respectively. The 3D Allen Brain Atlas was sourced from https://download.alleninstitute.org/informaticsarchive/allen_human_reference_atlas_3d_2020/version_1.

Since our study did not consider the hemispheric heterogeneity and given that the mouse brain atlas is symmetrical, we used the left hemisphere parcellation and applied mirror symmetry to generate the corresponding regions in the right hemisphere. This resulted in a human brain atlas in MNI space, containing 127 regions of interest (ROIs), including 105 cortical ROIs and 22 subcortical ROIs.

### Mouse atlas

The parcellation of the mouse brain was based on the Allen Mouse Brain Atlas^101^, accessed through the Allen Institute’s API. We aggregated this atlas using a hierarchical clustering approach by merging the labels of smaller brain regions into the larger brain region they belong to. This consolidation approach preserved anatomical relationships while reducing regional fragmentation. After excluding white matter and ventricular regions, we intersected the aggregated atlas with the spatial domains defined in our whole-brain spatial transcriptomic dataset^40^. The final mouse brain atlas in CCFv3 space comprised 66 regions of interest (ROIs), including 39 isocortical and 27 subcortical regions.

### Transcriptomic data processing AHBA data

The human gene expression data was obtained from the AHBA^32^. The data was downloaded from the Allen

Institute’s API and preprocessed using the *abagen* package^102^. During the preprocessing, we utilized data from six donors, each containing log2 expression values for 58,692 gene probes derived from multiple tissue samples. In parameter selection, we used *diff_stability* for probe selection. To preserve as many genes as possible, no restrictions were applied to intensity-based filtering. Additionally, to increase spatial coverage of the samples, we set *lr_mirror* to bidirectional. For each donor, we standardized gene expression both within samples and across samples using the scaled robust sigmoid method during gene expression matrix extraction. The *norm_matched* parameter was set to False. The *region_agg* parameter was set to None. All other parameters were left at their default settings. The gene expression matrix was obtained using the *get_expression_data* function from the *abagen* package. Based on the human brain atlas employed in our study, we obtained a gene expression matrix comprising 4,729 samples and 20,232 genes across all donors.

### Human single-nucleus data

The large-scale single-nucleus dataset for the human brain used in our study was provided by the Linnarsson laboratory^28^. This dataset includes single-nucleus transcriptomic data collected from multiple brain regions of three adult human donors. To minimize the differences caused by the sequencing process between individuals while ensuring a sufficient number of brain regions were included in the dataset, we assessed the variations in total UMI per cell (which can partially reflect batch effects) and the degree of overlap among sampled brain regions across donors. Ultimately, we retained the data from donors (H19.30.001 and H19.30.002) for further analysis. For the filtered single-nucleus dataset, we conducted preprocessing for each individual (*scanpy*, https://scanpy.readthedocs.io/en/stable), which mainly included the following steps:

(i) Cells with fewer than 200 detected genes and total UMI counts below 800 were removed. Additionally, genes expressed in fewer than 30 cells were filtered out. Subsequently, we normalized the UMI counts for each cell by scaling the total UMI counts to a target value of 10,000. To avoid the impact on the normalization process induced by disproportionately high expression of certain genes (e.g., ribosomal or mitochondrial genes), we excluded genes that accounted for more than 5% of the total expression in any single cell. After normalization, we applied a log2 transformation to the expression values of each gene.
(ii) We selected 78 common brain regions between the two donors (H19.30.001 and H19.30.002). Cell types with lower region-specific contributions, including ‘Choroid plexus’, ‘Ependymal’, ‘Miscellaneous’, ‘Splatter’, and ‘Vascular’, were excluded from the dataset. Furthermore, due to some inherent technical limitations in single-nucleus sequencing, such as the challenges in determining anatomical boundaries between donors and the unconstrained proportions of neurons and non-neurons in subcortical regions^28^, we evaluated the stability of cell type proportions across individuals in different brain regions, and filtered out samples or regions with low stability. Specifically, our approach is as follows: For each brain region, two to four samples were usually acquired from a single donor. First, we identified the cell types shared by both donors based on the ‘cluster’ annotation. Next, for each brain region, we calculated the pairwise Pearson *r* correlation coefficients between all samples from both donors using cell type proportions. The sample with the highest average similarity was defined as ‘regional reference’ and unstable samples with the correlation to reference below 0.6 were filtered out. Then, we recalculated the region-level similarity between donors and discarded regions with cross-donor correlations below the threshold (0.6) or those with more than a tenfold difference in cell counts. Finally, 55 out of the 78 brain regions with high stability were retained.
(iii) In order to maintain the independence of transcriptomic data between the two donors, we conducted total UMI regression separately within each donor. Furthermore, Considering the independence of data across donors and the non-dominant role of single-nucleus gene expression in data integration, we did not apply batch effect correction between donors.
(iv) To smooth regional gene expression, we employed a bootstrapping resampling method. Specifically, we randomly selected 100 cells within each region to compute the mean gene expression. This method is to better characterize transcriptional patterns at the regional level while ensuring effective integration with the AHBA microarray data. The sampling process repeated until the number of smoothed cells matched the original cell counts of that region.

### Mouse spatial transcriptomic data

The whole-brain ST array sequencing data from three adult male mice were preprocessed through steps including log2 transformation and cross-animal batch effect correction, followed by alignment to the CCFv3 standard space^40,103^. Each row of the processed gene expression matrix corresponds to a spot, and each spot is indexed by a unique Allen CCFv3 atlas label. We updated the spatial annotations according to the hierarchical relationships of different brain regions in the mouse brain atlas we used, ultimately resulting in a gene expression matrix with 20,023 spots and 15,239 genes.

### Integration of single-nucleus transcriptomic data with AHBA

The process of integrating single-nucleus data with AHBA to generate train data primarily consisted of three steps: alignment of the single-nucleus sampled brain regions with the Human Brain Atlas, selection of stable genes based on spatial patterns across datasets, and weighted combination of gene expression vectors.

(i) Alignment of the single-nucleus sampled brain regions with the AHBA: Since the sampled brain regions from human single-nucleus data do not correspond one-to-one with the Human Brain Atlas used in our study, we calculated the centroid distance between the ROIs of the two atlases in the standard MNI152 space, based on the optimal matching of distances and utilized the Brodmann Atlas^104^ as an intermediary to match the labels of the two datasets.
(ii) Transcriptomic data scaled: For the gene expression matrices of different modalities, we applied z-score normalization along the sample axis.
(iii) Selection of stable genes based on the similarity of spatial patterns across datasets: We integrated genes from the preprocessed single-nucleus gene expression matrix, the AHBA gene expression matrix, and the cross-species homologous genes (sourced from R *homologene* package) of the mouse spatial transcriptomic data, resulting in a total of 12,732 genes. Next, by aligning brain region labels between single-nucleus data and 3D Allen Brain Atlas, we acquired spatial locations of 42 single-nucleus sampled brain regions in MNI152 space. This alignment enabled us to utilize the AHBA data processing pipeline, as previously described, to compute average spatial expression patterns for each gene across single-nucleus sampled regions. Subsequently, we calculated the Pearson *r* correlation of each gene’s mean expression patterns across regions in both single-nucleus and AHBA data. Genes with cross-modal regional expression similarity greater than the median of overall similarity were retained. This step was conducted separately for the two donors in the preprocessed single-nucleus data. Finally, we identified the intersection of cross-modal stable genes in the two donors.
(iv) Weighted combination of gene expression vectors: A key strategy for data integration is that, for each sample of each brain region in the AHBA gene expression matrix, a specified number of smooth-cells are randomly sampled from the corresponding regions in the preprocessed single-nucleus data pool, and gene expression between the two datasets were weighted and averaged. In practice: we calculated the Pearson *r* correlation between the average gene expression of each brain region extracted from the AHBA and each smooth-cell from the corresponding brain region of preprocessed single-nucleus data. We retained the top 10% of smooth-cells with the highest correlation, which were then added to the random sampling pool. In the process of data integration, we recognized that although single-nucleus data offers a large-scale sample size, it has relatively low spatial resolution. We aimed to introduce a certain degree of variability in the AHBA using single-nucleus data to augment the training samples, while minimizing the excessive incorporation of single-nucleus information in each sample that could disrupt the gene expression heterogeneity across different regions. Hence, we employed relatively conservative parameters in this process. Specifically, in each random sampling, we selected 10 smooth-cells from the corresponding regions in the preprocessed single-nucleus data pool, and we weighted smooth-cell expression and AHBA sample expression in a ratio of 3:7.

We repeated the data integration process 100 times at both the cortical/subcortical scale and whole-brain scale, the resulting fused datasets served as raw input for the detached and shared deep neural network model to characterize region-specific transcriptional patterns.

### Capturing cross-species conserved transcriptomic patterns

#### Training deep neural network using integrated human transcriptomic data

To enhance the accuracy of anatomical region matching based on transcriptomic data between human and mice, we constructed a deep neural network to learn region-specific transcriptional patterns in the human brain^24^. Based on different transcriptomic data normalization strategies, we constructed two deep neural network models: detached model and shared model. The detached model consists of two independent parts: the cortical module and the subcortical module. Cortical model took separately normalized fusion data of the cortex as input, with an input layer dimension of 4,542 and an output layer of 105 cortical region labels. Subcortical model took separately normalized fusion data of the subcortex as input, with an input layer dimension of 5,063 and an output layer of 22 subcortical region labels. The shared model took fusion data normalized across the entire brain as input, with an input layer dimension of 5,451 and an output layer of 127 whole-brain region labels.

The integrated transcriptomic data was fed into the model after z-score normalization along the sample axis. All models utilized the rectified linear unit (ReLU) activation function during forward propagation, employed the cross-entropy loss function for loss calculation, and used the AdamW optimizer for weight updates with a weight decay set to 1×10^-6^. Number of hidden units: 500, 500, 500. The maximum learning rate during training was set to 6×10^-5^, with a learning rate optimization strategy of StepLR, and the maximum number of epochs for each model was 50. For each generated fused dataset, we repeated the model training 10 times, randomly dividing 10% of the data as a validation set during each training iteration, retaining the model parameters that performed best on the validation set. After training, for both AHBA and Mouse ST data, we fed the regional average expression matrices into the model and extracted the outputs from the final hidden layer, representing region-specific embeddings. Finally, we obtained the average embedding matrix from all repeated models and calculated the pairwise correlations between regions across species, which served as the foundation for constructing the homologous mapping relationships.

To validate the model’s generalization ability and the cross-subject stability of region-specific transcriptional patterns, we conducted cross-subject validation experiments. In detail, we generated independent training and testing datasets by randomly dividing the sample expression data from AHBA into two groups of three subjects each. We considered all independent combinations of the grouped AHBA subjects with two subjects of the single-nucleus data, integrating data for each combination as previously described. Finally, we evaluated the average confusion matrix on all independent test sets and the classification accuracy for each brain region.

These models were implemented in Python using *PyTorch* (https://pytorch.org) and the *skorch* package (https://skorch.readthedocs.io/en/stable).

### Gene enrichment analysis

We used average gene expression as input for gradient backpropagation to identify genes that most significantly contribute to region specificity. This process was implemented using the Integrated Gradients function from the *captum* python package (https://captum.ai). Next, for each brain region, we selected the top 20 genes with the highest weights to form a high-weight gene representative set. To establish the association between these high-weight genes and biological processes, we performed pathway and process GO enrichment analysis on the high-weight gene set. GO enrichment analysis was performed using the Metascape platform^105^ (https://metascape.org).

### Random walk-based graph representation learning Human DTI

We utilized diffusion-weighted imaging (DWI) data from the Human Connectome Project (HCP), which have been preprocessed following HCP’s standard pipeline (e.g., motion correction, eddy current correction, and gradient distortion correction). Further post-processing steps of tractography and connectome construction were carried out using MRtrix3 (http://mrtrix.readthedocs.io). Briefly, A tissue-segmented image was generated to enable anatomically constrained tractography (MRtrix command *5ttgen*). The multi-shell multi-tissue response function was estimated (MRtrix command *dwi2response msmt_5tt*), followed by multi-shell, multi-tissue constrained spherical deconvolution (MRtrix command *dwi2fod msmt_csd*). Subsequently, an initial whole-brain tractogram was generated with 2 million streamlines (MRtrix command *tckgen*), with a maximum tract length of 250 mm and a fractional anisotropy (FA) threshold of 0.06. The final set of streamlines was then parcellated into 127 brain regions according to our human brain atlas (MRtrix command *tck2connectome*).

### Mouse Tracer

The mouse connectome data was initially sourced from the Allen Mouse Connectivity Atlas^33^ (https://connectivity.brain-map.org) and subsequently processed by Coletta et al. using high-resolution models (100 μm³) released by Knox and colleagues^44,106^.

To derive the region-level connectivity matrix, we resampled the mouse brain atlas in our study to the voxel-scale connectome template. We then averaged all voxel-level connectivity values between each pair of brain regions according to index correspondence between the voxel connectivity matrix and the voxel-scale connectome template, resulting in a 66 ROIs connectivity matrix. Additionally, considering the differences in data scale between Tracer and DTI, we scaled the mouse Tracer connectivity matrix to align with the human DTI matrix as follow:

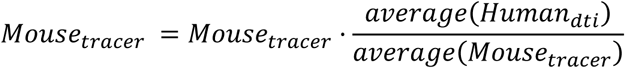

Without considering the directional projections between brain regions, we symmetrized the connectivity matrices of both human and mouse between left and right hemispheres, set the diagonal elements to zero, applied log2 transformation, and removed any negative values. Finally, we normalized each row of the matrix to complete the processing of the structural connectivity data.

### Embedding space generated from Human-Mouse graph

We used region-specific transcriptional similarity as weights to link human and mouse structural connectivity, constructing a Human-Mouse graph. Next, we pruned the links between human and mouse structural connectivity in the Human-Mouse graph based on the hierarchical brain relationships between species (**Supplementary Table.2**) and set the negative weight of links to 0, retaining only the links that connect nodes within the same hierarchy. To avoid introducing excessive prior information, we considered only major hierarchical brain correspondences between species to prune links. We also examined whether the region-specific transcriptional similarity patterns between species conformed to general brain organizational principles and removed mouse regions with clearly incorrect similarity, including secondary motor (MOs) and anteromedial visual (VISam), from the graph.

For the pruned Human-Mouse graph, we used the continuous bag of words Node2vec algorithm by Sporns et al. to construct graph embedding^56^. In this process, we repeated the model iteration 500 times. For each iteration, we set the number of walks per node to 800, with a step length of 40. The probability of random walks was weighted according to the edge weights in the graph. To analyze the impact of the walk strategy on graph embeddings, we set a gradient from breadth-first to depth-first search. We found that the recognition rank of potential homologous regions between human and mouse in the graph embeddings initially decreased as the walk leaned towards depth-first, then gradually increased. We selected the parameters (p=0.01 and q=0.1) that yielded the lowest recognition rank for potential homologous regions to construct the graph embeddings (**Supplementary Fig.10**). When constructing graph embedding, we set the window size to 5 and the dimensions to 40.

### Mapping whole-brain phenotypes comparison cross species

To enable quantitative comparison cross species, using the latent embedding we defined previously, we performed a dual regression method^23^ to map mouse imaging phenotype to human space (synthetic human brain). Briefly, the value of imaging phenotype in mouse brain of ROIs was first regressed by the mouse graph embedding to calculate a β matrix, then, we used the β matrix dot the graph embedding matrix of human, which will output an estimate vector consistent with the ROIs number of the human brain. This process was repeated 500 times, and the final estimate vector was the average of the 500 results. The same procedure was applied in reverse for human-to-mouse mapping.

### Mapping Macroscopic imaging phenotypes cross species Human resting-state fMRI

The resting-state fMRI data were obtained from the HCP Young Adult 1200 Subjects Data Release, consisting of 998 participants (531 females, aged 22-37 years, mean age: 28.7 ± 3.7 years). Resting-state fMRI scans were acquired across two sessions (acquired on different days) using a customized 3T Siemens Skyra scanner. Each session included two 15-minute runs with opposite phase encoding directions (left-right and right-left), with each run consisting of 1,200 timepoints. The scans were conducted with the following parameters: TR = 720 ms, TE = 33.1 ms, and 2mm isotropic voxels. The functional MRI data were registered to the standard MNI152 space, and processed using ICA-FIX^107^ to remove motion artifacts and other noise components. The detailed information can be found at https://www.humanconnectome.org/study/hcp-young-adult/data-releases.

### Mouse resting-state fMRI

The resting-state fMRI data for awake mice were made publicly available by Gozzi et al^58^. Resting-state fMRI data were obtained from 10 adult male mice (aged < 6 months) using a 7T Bruker MRI scanner. Each mouse underwent a long-duration scan consisting of one run with 1920 time points. Each run was scanned using a single-shot echo planar imaging (EPI) sequence with the following parameters: TR = 1000 ms, TE = 15 ms, and 18 coronal slices (voxel size: 230 × 230 × 600 μm).

Preprocessing of fMRI data was carried out utilizing the AFNI, FSL, and ANTs software packages following several previous work^10,58,108^. Specifically, we excluded the first 2 minutes of the time series to avoid interference from machine noise and initial acquisition instability. Subsequently, the resting-state fMRI data underwent time despiking (*3dDespike*, AFNI) and motion correction (*3dvolreg*, AFNI), resulting in an in-house fMRI mean template with a spatial resolution of 0.23 × 0.23 × 0.6 mm³ (*fslmaths*, FSL). We manually skull-stripped each in-house average template and applied this to the original fMRI images. The skull-stripped fMRI data were then registered to the in-house mouse brain template (*antsRegistration*, ANTs). Following previous denoising methods^58^, we regressed out the average cerebrospinal fluid signal along with 24 motion-related covariates, including three translational and rotational parameters estimated during motion correction, their temporal derivatives, and corresponding squared values. The time series were then detrended and bandpass filtered (0.01-0.1 Hz). Global signal regression was not applied^95^. Additionally, we performed frame-wise head motion correction on the time series, removing time points with framewise displacement (FD) greater than 0.065. Finally, we applied Gaussian spatial smoothing (kernel size=0.5 mm) to complete the preprocessing of mouse fMRI.

### Macroscopic imaging phenotypes analysis

We characterized the macroscopic functional organization of the mouse brain using functional gradients, temporal co-activation patterns (CAPs), and seed-based functional connectivity. Human and mouse functional gradients used in this study were previously published^41,44^. Following Gozzi et al^58,62^, CAPs and seed-based functional connectivity were computed from the preprocessed resting-state fMRI data of awake mice.

For mouse CAPs, we conducted PCA dimensionality reduction on the group-level time series matrix from all frames of the 10 mice along the voxel dimension. Next, we performed k-means clustering on the dimensionality-reduced time series × PCs matrix. We set the number of clusters to 6, used spatial correlation as the distance metric, a maximum of 500 iterations, and initialized with 5 different random seeds. After clustering, we averaged the PCs belonging to the same category to reconstruct the original 3D brain region activation patterns. Finally, we calculated the Pearson *r* correlation coefficients between different cluster centers as a measure of topological similarity, averaging the CAP and corresponding negative anti-CAP with the highest negative correlation coefficient.

In the mouse seed-based functional connectivity analysis, we set the size of the seed points to 4×4×4 voxels. These seed points comprised: the prefrontal cortex (PFC) and retrosplenial cortex (RS) from the default mode network (DMN); the insula (Ins) from the salience network (SN); the primary motor region (M1) from the motor network; the primary visual region (VISp) from the visual network. We extracted the average time series for each seed point and performed Pearson *r* correlation with the whole-brain voxel time series. The correlation maps for individual subjects were averaged across all subjects after applying Fisher’s r-to-z transformation. Next, we set the significant voxel (two-tailed t-test, *P*<0.01, FDR corrected) within each network to 1 and the remaining voxels to 0, generating a binarized network mask.

The cortical 7 networks in human and functional gradients were provided by Yeo et al. ^60^ and Smallwood et al.^41^, respectively. For human CAPs, considering the high computational cost of clustering in the original voxel space, we generated a map of 10k ROIs to extract time series. This approach not only reduces computational complexity but also preserves voxel-level information to a large extent. The process for generating the 10k ROIs atlas can be referenced from https://github.com/tyzhang97/Fine-grained-Brain-Atlas/tree/main. Next, we utilized the preprocessed fMRI data as described above to extract the group-level time series matrix for all frames from all subjects. Similarly, we performed PCA dimensionality reduction on the time series matrix along the ROIs dimension (PCs = 1000, interpretability of variance = 0.659) and executed k-means clustering on the resulting time series x PCs matrix. We set the number of clusters to 8, while keeping the other parameters consistent with those used in the mouse CAPs computation. For the obtained clustering results, as described in the identification of mouse CAPs, we averaged the PCs belonging to the same category to reconstruct the original 3D brain region activation patterns. Subsequently, we calculated the Pearson *r* correlation coefficients between different cluster centers as a measure of topological similarity, averaging the CAPs and anti-CAPs with the highest negative correlation coefficients.

The macroscopic network maps for both human and mouse were aligned to the atlas used in our study. As the mouse functional gradients and Yeo 7 networks include only cortical regions, we retained only cortical measurements in subsequent analyses, setting subcortical values to zero in all macroscopic networks. Finally, for the functional gradients and CAPs, we extracted regional activation value of each ROI from the atlas. For the seed masks and Yeo 7 networks, we calculated the ratio of network voxel counts to the total voxel counts for each ROI to assess the region’s involvement in the network.

### Optogenetic fMRI

Processed optogenetic fMRI data for the insula and dorsal raphe nucleus (DRN) were made available by Grandjean et al.^23,109^. We downloaded all the z-statistic maps of insula stimulation and DRN stimulated with 20 Hz, 5 ms pulse width without drug intervention. We subtracted the control group’s average z-statistic map from the experimental groups to obtain whole-brain activation patterns for the insula and DRN. All maps were aligned to the atlas used in our study, from which we extracted the average activation intensity of each ROI. Then, we created a virtual optogenetic activation map of the human brain based on our proposed framework.

To establish the relationship between optogenetic modulation and behavior, we employed Neurosynth (https://neurosynth.org) in the generated human optogenetic activation map. Neurosynth is a meta-analytical platform capable of generating keyword-based statistical functional maps that display activation patterns of specific cognitive processes or psychological terms^8^. We retained the top 20 brain ROIs with the highest average activation intensity of the generated human optogenetic activation map. We then calculated the overlapping voxels between these ROIs and the term-based statistical maps from Neurosynth (normalized by all activation voxel counts of term-based statistical maps) to assess the potential behavioral and cognitive functions regulated by the optogenetic activation circuits. To visually articulate the results, we employed a word cloud to depict the foremost 15 terms associated with two synthetic optogenetic maps, respectively.

### Deformation-based morphology analysis in autism across species

#### Deformation-based morphology

All mice used in this study were from a large set of autism mouse models, composed of separate cohorts from various laboratories, and sent to the Mouse Imaging Centre in Toronto for neuroimaging^13^. We filtered out missing genotypes and mutations related to the Y chromosome, ultimately retaining data on mutations involving 13 different genes.

We employed the deformation-based morphometry (DBM) method to quantify regional volumes in the mouse structural imaging data. All images were corrected for intensity nonuniformities and then aligned to the unbiased global average template provided by the dataset. The registration process was performed using ANTs^110^, involving cascaded linear (a 6-parameter followed by a 12-parameter) and nonlinear stages. After concatenating all derived transformations into a composite deformation field, we can compute the Jacobian determinant map defined over the template, which encodes the local volume of each voxel. The ABIDE I data had been processed by fMRIPrep^111^ anatomical workflow. Specifically, T1w images were bias-corrected, skull-stripped, and registered to the MNI152 template using ANTs. Analogous to the DBM analysis performed for mouse data, we calculated Jacobian maps based on the deformation fields from individual space to template space and then we can measure the regional volume. We registered our atlas to the corresponding template in the deformation-based analysis, ROI volumes for human and mouse brain were calculated by summing Jacobian values within each region and were normalized by the mean ROI volume to eliminate the influence of global volume.

#### Normative model

After obtaining ROI volumes for individuals with autism and controls, we applied a normative model to characterize abnormal regional volume deviations in autism. Given the heterogeneity and sample size differences across sites in the ABIDE I dataset, we used 468 participants from four sites where the number of autism subjects was greater than 40. The selected sites are: (1) NYU, with 79 ASD and 105 controls; (2) USM, with 58 ASD and 43 controls; (3) UM_1, with 55 ASD and 55 controls; and (4) UCLA_1, with 41 ASD and 32 controls. Next, we developed a normative model by training a Gaussian Process Regression on the selected sites from ABIDE I controls, using age, gender, and sites as covariates. This process was implemented using the PCNtoolkit^71^ (https://pcntoolkit.readthedocs.io). We quantified the deviation patterns of autism from the normative field established based on neurotypical subjects, as follows:

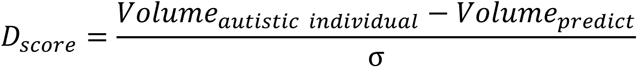

where Volum*e*_autistic individual_ represents the initial ROI volume of autism,*Volume*_predict_ is the expected ROI volume estimated from the Gaussian Process Regression, and σ is the square root of variance estimated from the GPR. Subsequently, we applied a general linear model to regress out the effects of age, gender, and site on the individual-level deviation patterns.

#### Morphological differences in mouse autism model

We first assessed age and sex matching between autism model mice and wild-type controls for each genotype, confirming no significant differences. Referring to the method described by Ellegood et al. ^13^and considering the limited number of mice within each genotype, we did not regress out age and sex within each genotype. For autistic-normal comparisons, we calculated Cohen’s d scores for each brain region between autistic and normal mice within each genotype, and we statistically evaluated the number of regions with significant differences (two-tailed test, *t* > 1.96). For subsequent analyses, we excluded genotypes with fewer than 15 significantly altered regions due to their limited statistical strength. The remaining models included: *dvl1^-/-^* (n=55), *magel2^-/-^* (n=24), *slc6a4^-/-^* (n=22), *sgsh^-/-^* (n=21) and *shank3^-/-^*(n=16). The volumetric deviation patterns of these mouse models were mapped to human brain space using our TransBrain framework and subsequently correlated with volumetric alterations observed in human autism data.

## Data availability

The Allen Human Brain Atlas is available at http://human.brain-map.org, the single-nucleus dataset for the human brain is available at https://storage.cloud.google.com/linnarsson-lab-human. The processed mouse gene expression data and 3D coordinates of spots can be accessed at https://www.molecularatlas.org. The neuroimaging data for this research were procured from the publicly available Human Connectome Project Young Adult (HCP-YA) Project, which can be found at https://db.humanconnectome.org. The awake mouse resting fMRI dataset can be downloaded from https://data.mendeley.com/datasets/np2fx99hn6/2. Processed optogenetic fMRI datasets for the insula and dorsal raphe nucleus are available at https://doi.org/10.34973/raa0-5z29. The preprocessed ABIDE I data of 1102 subjects is available at https://fcon_1000.projects.nitrc.org/indi/abide. The mouse structural imaging dataset related to autism is available at https://www.braincode.ca/content/public-data-releases#dr001. Other atlases and imaging datasets used in this study were all obtained from previous publications, as detailed in the Methods section.

## Acknowledgements

This work was supported by the Science and Technology Innovation 2030-Brain Science and Brain-inspired Intelligence Project of China (2022ZD0211900, 2022ZD0207700, and 2021ZD0200102), the National Natural Science Foundation of China (NSFC) (32192411, 32122037, 31971028, and 32222032), the New Cornerstone Science Foundation, Changping Laboratory, and the Open Research Fund of the State Key Laboratory of Cognitive Neuroscience and Learning.

## Author contributions

A.L. and X.W. led the project. A.L., X.W. and S.H. were responsible for the study concept and the design of the study. S.H., T.Z. and A.L. analyzed the data. S.H., A.L. and X.W. created the figures and wrote the paper. T.Z., C.D., Y.S. and Y.P. aided in the writing of the paper. X.L., K.L., Q.W., Y.H., F.X., X.T., J.X., C.Z., Y.Z., L.L. and B.L. participated in discussions of the results and the paper.

